# Kinetic basis for DNA target specificity of CRISPR-Cas12a

**DOI:** 10.1101/355917

**Authors:** Isabel Strohkendl, Fatema A. Saifuddin, James R. Rybarski, Ilya J. Finkelstein, Rick Russell

## Abstract

Class II CRISPR-Cas nucleases are programmable via a single guide RNA, enabling genome editing applications in nearly all organisms. However, DNA cleavage at off-target sites that resemble the target sequence is a pervasive problem that remains poorly understood mechanistically. Here, we use quantitative kinetics to dissect the reaction steps of DNA targeting by *Acidaminococcus sp* Cas12a (also known as Cpf1). We show that Cas12a binds DNA tightly in two kinetically-separable steps. Protospacer-adjacent motif (PAM) recognition is followed by rate-limiting R-loop propagation, leading to inevitable DNA cleavage of both strands. Despite the functionally irreversible binding, Cas12a discriminates strongly against mismatches along most of the DNA target sequence, implying substantial reversibility during R-loop formation –a late transition state– and the absence of a ‘seed’ region. Our results provide a quantitative underpinning for the DNA cleavage patterns measured in *vivo* and observations of greater reported target specificity of Cas12a than the Cas9 nuclease.

## INTRODUCTION

CRISPR-Cas systems have emerged as revolutionary new tools for gene editing applications (Hsu et al., 2014). The nuclease enzymes in these systems can be readily programmed by the rational design of a CRISPR RNA (crRNA) sequence. The RNA-guided nucleases identify potential targets via Protospacer-adjacent motif (PAM) recognition and then form an R-loop between the guide RNA and the complementary target to license DNA cleavage. The information inherent in the ~20 base pairs formed by the crRNA and the target DNA is sufficient, in principle, to uniquely target a single sequence, even in large eukaryotic genomes.

Class II CRISPR-Cas systems are particularly useful because they use a single polypeptide for both target recognition and cleavage of the target DNA or RNA. However, in eukaryotic cells the prototypical Class II enzyme, Cas9, invariably falls short of the goal of a single unique site, as related ‘off-target’ sequences are also recognized and cleaved at significant levels (Sternberg and Doudna, 2015). This existence of off-target cleavage events is not surprising, as molecular interactions are never perfectly specific. For example, mismatches in DNA/RNA duplexes typically incur penalties of only 2-5 kcal/mol in solution (Sugimoto et al., 1995; Watkins et al., 2011). Even with one or two mismatches, a 20-base-pair helix is stable (Herschlag, 1991) and complexes formed with imperfect targets may be sufficiently long-lived to favor cleavage over enzyme dissociation (Bisaria et al., 2017). Despite the importance of these off-target cleavage events, we still lack a full understanding of the biophysical basis of the specificity of CRISPR-Cas enzymes.

Cas12a (also known as Cpf1) is a Class II enzyme (Makarova et al., 2015; Shmakov et al., 2015; Zetsche et al., 2015) that has recently emerged as a more specific alternative to Cas9 (Figure 1A) (Kim et al., 2016; Kleinstiver et al., 2016). Indeed, *in vivo* editing was achieved in mammalian cells (Toth et al., 2016; Tu et al., 2017; Zhong et al., 2017), and both *Acidaminococcus sp*. BV3L6 Cas12a (AsCas12a) and *Lachnospiraceae* bacterium Cas12a (LbCas12a) showed little or no tolerance for mismatches (Kim et al., 2016; Kim et al., 2017; Kleinstiver et al., 2016). In contrast, Cas9 discriminates strongly against mismatches only within the first ~10 base pairs proximal to the PAM and tolerates PAM-distal mismatches. Cas12a is also capable of processing its own precursor crRNA, unlike Cas9, and can therefore be used for multiplexed applications (Fonfara et al., 2016; Zetsche et al., 2017).

One striking question that arises is how different CRISPR-Cas enzymes can have different levels of specificity despite using the chemically identical R-loop as the source of sequence-specific DNA binding. A common feature of CRISPR-Cas systems and other RNA-directed enzymes is that the targeting RNA includes a ‘seed’ region. The seed region, which for CRISPR-Cas enzymes is a subset of nucleotides within the crRNA that base pairs with PAM-proximal nucleotides, is highly sensitive against mismatches and therefore contributes importantly to target affinity and specificity (Gorski et al., 2017; Jinek et al., 2012; Wiedenheft et al., 2011). A crystal structure of Cas12a from *Francisella novicidia* (FnCas12a) showed that the bound crRNA includes a 5-nucleotide region that is pre-ordered for base pair formation (Swarts et al., 2017) with PAM-proximal nucleotides. This pre-ordered crRNA, however, cannot account for the high cleavage specificity seen throughout the R-loop beyond these five nucleotides.

**Figure 1.**
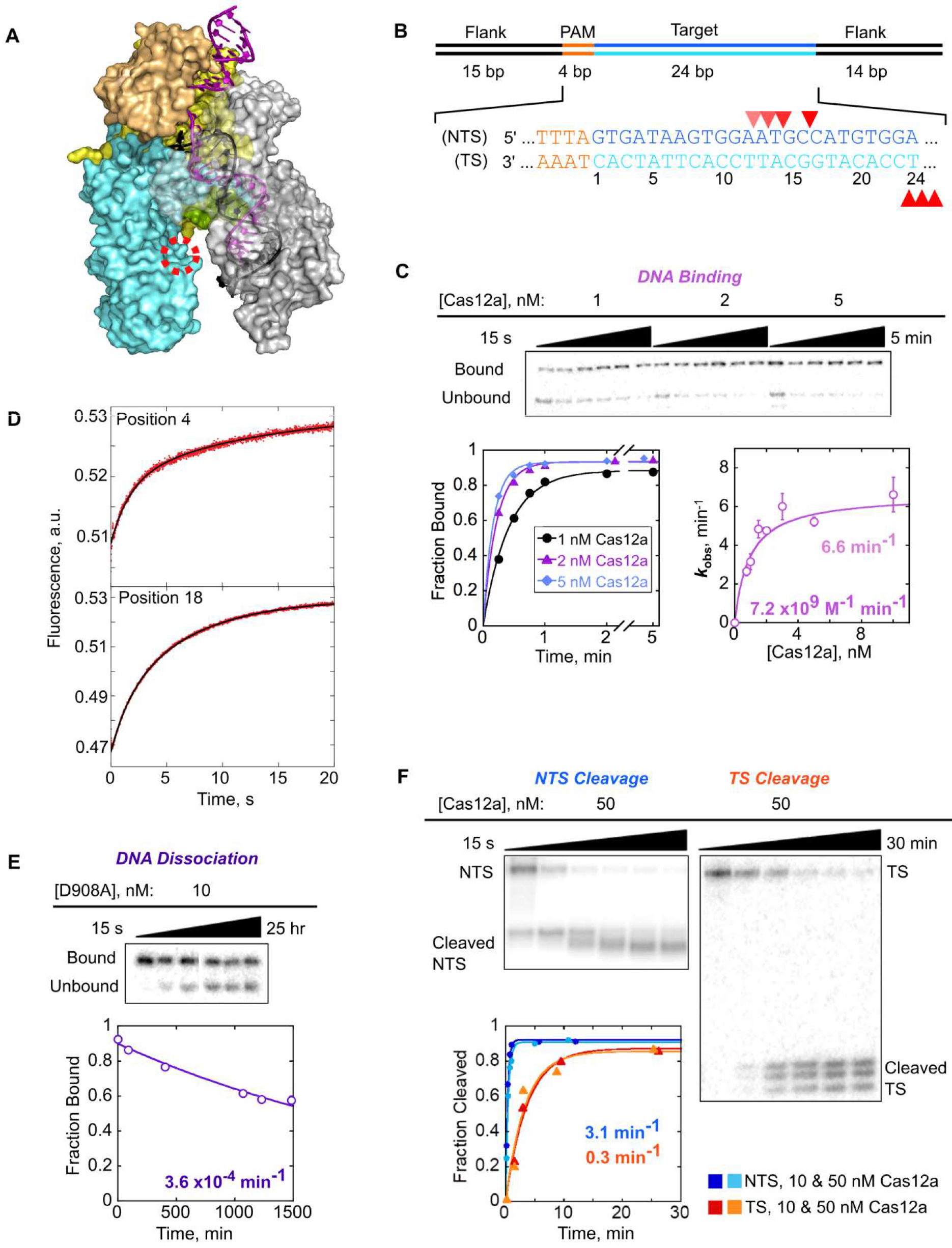
Kinetics of Cas12a binding and cleaving a matched target DNA. (A) Crystal structure of AsCas12a in complex with crRNA (black) and DNA (purple) (PDB: 5b43). Nuclease Lobe; grey: Recognition Lobe; Red, dashed circle: RuvC nuclease active site where the D908A mutation is located. The 20-bp R-loop formed between the crRNA and the TS lines the Recognition Lobe, and the NTS is predicted to line the Nuclease Lobe. One of the REC domains is transparent to increase visibility of the R-loop. (B) Schematic and sequence of the target DNA substrate. Red triangles mark cleavage sites. Lighter triangles on the NTS indicate cleavage sites in ‘trimming’ events that follow the initial cleavage event. (C) Representative gels and plots showing time dependences of Cas12a binding to a matched DNA target, and a corresponding plot of the observed rate constant as a function of Cas12a concentration. Error bars reflect the SEM of 3-5 replicates. See also Figure S1. (D) 2-aminopurine (2-AP) fluorescence curves showing R-loop propagation for wild-type Cas12a in the presence of EDTA after rapid binding to DNA. 2-AP was substituted at position 4 or 18 of the NTS as indicated. See also Figure S2. (E) Representative time course gel and plot showing dissociation of Cas12a(D908A) from the matched target DNA. (F) Representative gels showing multiple product bands upon Cas12a-mediated cleavage of the NTS (left) and TS (right). Reactions included 10 nM or 50 nM Cas12a as indicated.

Despite the many published crystal structures (Dong et al., 2016; Gao et al., 2016; Swarts et al., 2017; Yamano et al., 2016; Yamano et al., 2017) and *in vivo* targeting results, little is known about how Cas12a achieves its high specificity.

Here, we report a quantitative dissection of the binding and cleavage reactions of Cas12a, pre-loaded with a defined crRNA, toward a perfectly matched DNA target and a series of targets with single mismatches. We find that DNA binding is rate limiting for cleavage, both for matched and mismatched targets. Significant discrimination against mismatches is observed across most of the R-loop, suggesting reversibility within the process of R-loop formation itself and the absence of a defined seed region. Our results provide fundamental insights into DNA targeting by Cas12a and the origins of greater DNA target specificity of Cas12a than Cas9. In addition, our work suggests strategies for further improvement of specificity through enzyme engineering.

## RESULTS

### A kinetic framework for recognition and cleavage of a matched DNA target

We first generated a complete kinetic framework for Cas12a on a matched DNA target. The rates of DNA binding and dissociation, as well as cleavage of each DNA strand, were measured using a crRNA-loaded AsCas12a (henceforth referred to as Cas12a) and an oligonucleotide DNA target (Figure 1B). Our target DNA included a 5´;-TTTA PAM sequence, which conforms to the consensus 5´-TTTV-3´ PAM (Zetsche et al., 2015), adjacent to 24 base pairs that matched the crRNA targeting sequence. The target DNA was ^32^P labeled at the 5´ end of either the target strand (TS) or the non-target strand (NTS).

The second-order rate constant for Cas12a binding (*k*_on_) was determined by incubating various concentrations of Cas12a with a trace amount of NTS-labeled duplex. Reactions aliquots were quenched at various times by adding an excess of the same unlabeled duplex and analyzed by native polyacrylamide gel electrophoresis (Figure 1C). The dependence of the observed rate constant on Cas12a concentration gave a *k*_on_ value of 7.2 (± 0.2) × 10^9^ M^−1^ min^−1^, within 1-2 orders of magnitude of the macromolecular diffusion limit. The same association rate constant was observed with the nuclease-inactive Cas12a(D908A) (Yamano et al., 2016; Zetsche et al., 2015), indicating that cleavage of the DNA does not impact the observed binding rate (Figure S1). A hyperbolic dependence with an apparent plateau was observed for both Cas12a and Cas12a(D908A), indicating that binding includes two detectable steps. We infer that the *K*_1/2_ value of 0.9 ± 0.2 nM reflects the equilibrium constant for an initial complex, most likely resulting from PAM recognition (Gong et al., 2018). The maximal rate constant (*k*_max_) of 6.6 ± 0.4 min^−1^ reflects that a first-order step following initial binding, which we show below to be R-loop formation, limits the overall binding rate at high Cas12a concentrations. Cas12a(D908A) behaved similarly but with a somewhat larger *k*_max_ value 15 ± 4 min^−1^, suggesting that the mutated residue within the RuvC domain may be involved in R-loop formation.

To directly measure R-loop formation, we used stopped-flow fluorescence with a DNA substrate that included a 2-aminopurine (2-AP) at either position 4 or 18 of the NTS to report on the early and late stages of R-loop propagation (Figure S2A). The fluorescence of 2-AP is strongly quenched by stacking when it is base paired, making 2-AP a sensitive probe for monitoring base-pairing changes in Cas9 (Gong et al., 2018) and many other nucleic acid systems (Jones and Neely, 2015; Raney et al., 1994; Russell et al., 2006; Tamulaitis et al., 2007). Potential complications associated with DNA cleavage were avoided by adding EDTA to chelate Mg^2+^ or by using Cas12a(D908A) in the presence of Mg^2+^. In the presence of EDTA, the 2-AP fluorescence signal increased with a rate constant of 6 ± 2 min^−1^ for wild-type Cas12a (Figure 1D). The rate constant was not affected by the position of the 2-AP probe and statistically indistinguishable from the plateau observed in the binding kinetics experiments above (Figure 1C). Cas12a(D908A) in the presence of Mg^2+^ gave a similar fluorescence increase and a somewhat larger rate constant of 10 ± 1 min^−1^, which was unaffected by the position of the 2-AP or the presence of Mg^2+^ and was indistinguishable from the *k*_max_ value observed in DNA binding experiments (Figure S2B). All of the fluorescence traces also included a more rapid transition that was best modeled as a decrease in 2-AP fluorescence following the increase from R-loop formation (Figure 1D and Figure S2). This transition may reflect a conformational change that follows R-loop formation. Together, our results indicate that R-loop formation by Cas12a occurs with a rate constant of ~6 min^−1^. The independence of the rate constant on the 2-AP position indicates that, upon initiation, the process of R-loop propagation is fast and largely complete within a few seconds.

The rate constant for dissociation of Cas12a(D908A) from the target DNA (*k*_off_, Figure 1E) was measured using pulse-chase experiments and native gel separation. Dissociation was extremely slow in the presence of Mg^2+^, with a *k*_off_ value of 3.6 (± 0.2) × 10^−4^ min^−1^ (t_1/2_ = 32 ± 2 hrs). As a control, we compared the wild-type Cas12a and Cas12a(D908A) in the presence of EDTA to block DNA cleavage. The *k*_off_ values were 0.015 (± 0.002) and 0.014 (± 0.002) min^−1^ for Cas12a and Cas12a(D908A), respectively (Figure S1B), indicating that the D908A substitution does not affect the lifetime of the complex. In addition, Mg^2+^ increases the lifetime of the ternary complex by 40-fold, perhaps by favoring the conformational transition that follows R-loop formation or by contributing more directly to R-loop stability in the complex.

Our measured rate constants give a calculated *K*_d_ value of 49 ± 6 fM (*k*_off_ / *k*_on_). This value is 10^4^-fold lower than the recently reported value of 0.1 nM from single molecule fluorescence experiments (Singh et al., 2017) and 10^6^-fold lower than that determined via binding affinity assays with FnCas12a in the absence of Mg^2+^ ion (Fonfara et al., 2016). These previous studies used direct measurements of the fraction of DNA target bound rather than kinetics measurements, and they did not provide the >32 hr incubation required to reach equilibrium, presumably contributing to the apparently weaker binding of Cas12a. Although our measured equilibrium constant is extraordinarily low for an enzyme-substrate interaction, reflecting a very stable complex, the value is orders of magnitude higher than the predicted *K*_d_ for a 20 base pair RNA:DNA duplex of the same sequence (~0.02 fM) (Sugimoto et al., 1995).

We next measured cleavage of the non-target strand and the target strand by individually radiolabeling each strand in separate reactions (Figure 1F). For each strand, hyberbolic fits to the observed rate constants from multiple Cas12a concentrations were used to determine the rate constants for cleavage (*k*_c_^NTS^ and *k*_c_^TS^). We obtained a maximal rate constant of 3.1 ± 0.4 min^−1^ for NTS cleavage and a 10-fold lower value for TS cleavage (0.30 ± 0.01 min^−1^). The slower TS cleavage is consistent with a recent report that the NTS is expected to line the Nuclease Lobe of Cas12a, which contains the RuvC nuclease domain (Stella et al., 2017; Swarts et al., 2017) (Figure 1A), and is thus poised for cleavage. In contrast, the R-loop TS:crRNA heteroduplex lines the Recognition Lobe and may require a substantial conformational change to position the TS for cleavage.

Close inspection of the DNA cleavage reactions revealed multiple products for each strand (Figure 1F). For the NTS, the initial cleavage event occurred at position 16 and was followed by apparent trimming of up to 4 nucleotides, with a rate constant of 0.72 ± 0.06 min^−1^, resulting in accumulation of faster-migrating products. For the TS, multiple cleavage products appeared with the same time dependence and maintained their relative intensities over time. The apparently imprecise cleavage, occurring at positions 23-25, suggests flexibility of this region of the TS. The lack of further cleavage events on the TS most likely reflects rapid dissociation of the PAM-distal product, which includes the radiolabeled end of the TS, after both DNA strands have been cleaved (Singh et al., 2017). Previous studies of FnCas12a have differed in the reported positions of DNA cleavage (Fonfara et al., 2016; Stella et al., 2017; Zetsche et al., 2015). Our findings of trimming and imprecision in the positions of DNA cleavage reconcile these previous results and have biological implications for the processing of the DNA ends (see Discussion).

Together, our results show that DNA cleavage by Cas12a occurs orders of magnitude faster than dissociation from a matched DNA target, indicating that essentially every complete binding event of Cas12a results in DNA cleavage (Bisaria et al., 2017). A key consequence of this irreversible binding is that the specificity of Cas12a against mismatches must be determined by the relative binding kinetics, not by differences in equilibrium binding.

### Single mismatches throughout the R-loop slow Cas12a binding and suggest the absence of a ‘seed’ region

To measure the specificity of Cas12a against mismatches between the crRNA and target DNA, we introduced single base pair changes in the DNA. Mismatches were chosen throughout the R-loop and were generated by inverting the nucleotides of the NTS and TS (except at position 14, where this change would have created an undesired PAM sequence). The *k*_on_ values were reduced substantially for mismatched DNA targets, with pronounced effects along nearly the entire length of the R-loop (Figure 2, Figure S3, Table S2). PAM-proximal mismatches gave ~150– to 600-fold decreases and PAM-distal mismatches gave ~25– to 80-fold decreases. Additionally, we found that the identity of the nucleotide mismatch can impact the binding rate. We were intrigued that C16G slowed binding by nearly 70-fold, despite being distal to the PAM, so we tested a second mismatch, C16T, which slowed binding by only 3-fold. A mismatch at position 21 did not decrease the rate of binding, supporting previous findings that a conserved aromatic residue (W382) stacks on position 20 of the RNA-DNA duplex and prevents further R-loop propagation (Swarts et al., 2017; Yamano et al., 2016).

The sensitivity of Cas12a to mismatches indicates that all positions tested (out to position 18) contribute to the binding rate constant. This result was surprising because when a RNA-DNA duplex forms in the absence of a protein co-factor, the transition state is reached after formation of only a few base pairs (Li et al., 2006; Porschke, 1977; Woodside et al., 2006b). Thus, only these base pairs contribute to the rate and specificity of duplex formation (Woodside et al., 2006a). In contrast, our results indicate that when Cas12a-crRNA binds to its target DNA, formation of the R-loop has a later transition state, with more base pairs formed. Indeed, assuming that the R-loop begins to form adjacent to the PAM sequence (Sternberg et al., 2014; Szczelkun et al., 2014), the sensitivity of the binding rate to PAM-distal mismatches implies that propagation of the R-loop remains readily reversible until nearly the entire loop is formed. Biologically, this late transition state would be expected to permit Cas12a to discriminate against mismatches in spite of the irreversible nature of the overall binding process.

**Figure 2.**
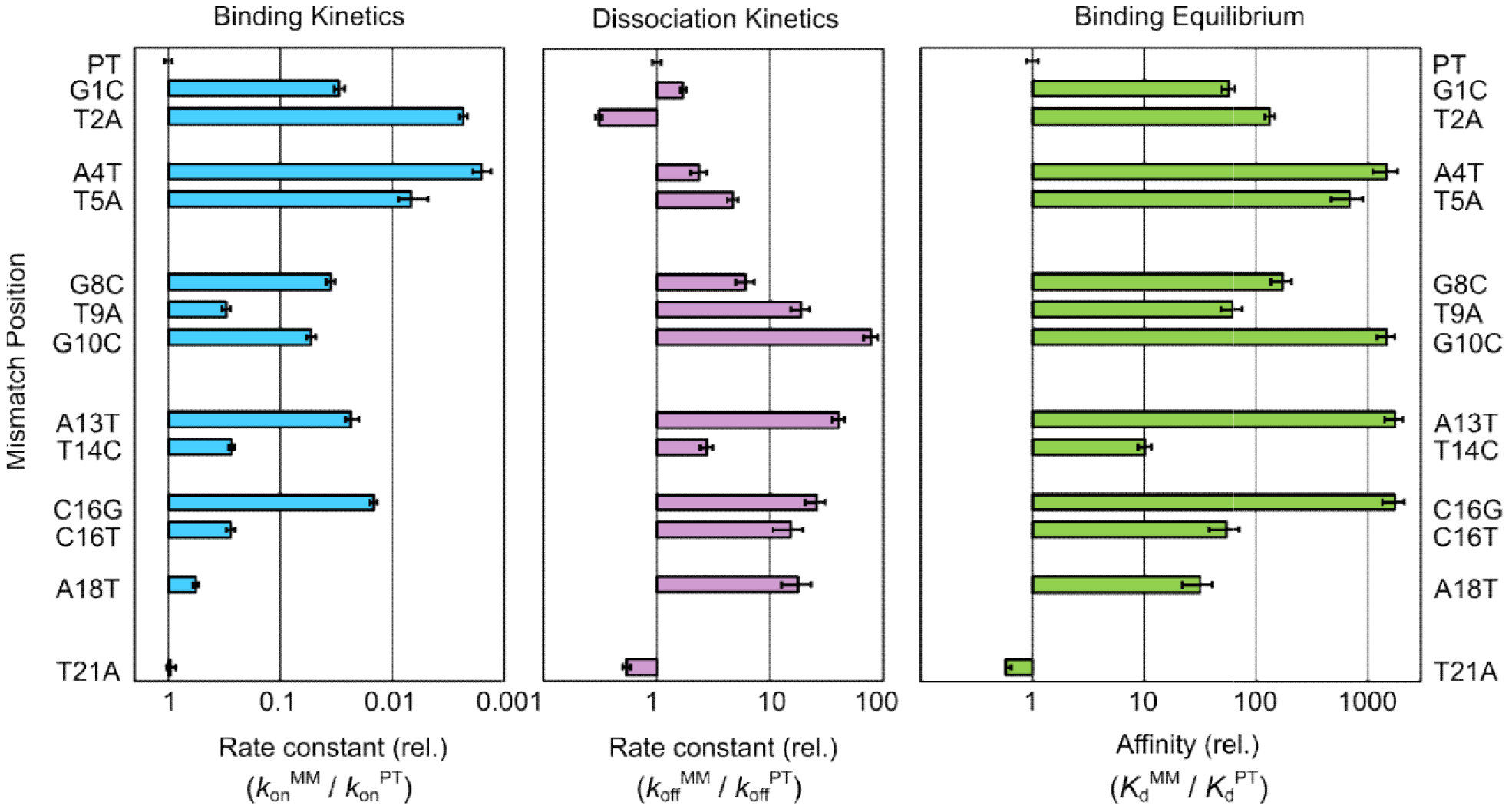
Mismatches throughout the R-loop reduce the affinity of Cas12a for target DNA. Bar graphs, left and center, represent the rates of binding and dissociation of Cas12a for a mismatch target (MM) normalized by the corresponding rates for the matched (*i.e.* perfect) target DNA (PT). The combination of the relative rate constants gives the overall effect on affinity, as depicted in the graph to the right. Error bars represent the normalized standard error of the fit and the normalized SEM (n ≥3) for binding and dissociation, respectively. Target names are defined as the mismatch position and substituted nucleotide within the NTS. See Also Figure S3 and Table S1 and S2.

To further explore the effects of sequence mismatches on Cas12a binding, we measured Cas12a dissociation from this series of DNA targets. All but one mismatch within the R-loop increased the dissociation rate (Figure 2), as is expected for destabilizing changes to the bound complex. Intriguingly, the largest increases in the dissociation rate were caused by PAM-distal mismatches. This effect has also been observed for Cas9 (Boyle et al., 2017; Szczelkun et al., 2014). PAM-distal mismatches increased the rate of dissociation 6– to 80-fold, except one substitution that gave a smaller increase of 3-fold (T14C), perhaps because the mismatch can form a dG•rU wobble base pair. In contrast, PAM-proximal mismatches increased the rate of Cas12a dissociation only up to 5-fold. The larger effects of PAM-distal mismatches on dissociation support the late transition state model proposed from our *k*_on_ results.

The combined effects of mismatches on *k*_on_ and *k*_off_ reveal a high level of thermodynamic discrimination by Cas12a for its perfect target DNA (Figure 2). Mismatches throughout the R-loop weaken binding substantially, with typical effects ranging from 50-fold to nearly 1000-fold. Although the effects of single mismatches are large, the resulting complexes are still quite stable, with sub-nanomolar affinity. The thermodynamic effects of mismatches do not display a detectable trend across the R-loop, providing evidence against a seed region of enhanced discrimination against mismatches in equilibrium binding.

### Mismatches do not impact DNA cleavage rates

We next measured cleavage of both strands of the mismatched targets (*k*_c_^NTS^ and *k*_c_^TS^) by using labeled targets as above and saturating Cas12a concentrations (Figure 3A). Mismatches decreased *k*_c_^NTS^ in a manner that paralleled the slower R-loop formation determined from our binding data (Figure 3B). The correspondence of these rate constants across a range of >100-fold suggests that the rate of NTS cleavage is limited by propagation of the R-loop.

Because TS cleavage is approximately 10-fold slower than NTS cleavage for the matched target, we expected that mismatches that slow R-loop propagation, and thus NTS cleavage, by less than 10-fold would not decrease the observed rate of TS cleavage. For mismatches that give larger effects, cleavage of both strands would be rate-limited by R-loop propagation, such that the observed rate of TS cleavage would be decreased to that of R-loop propagation and NTS cleavage (modeled by solid line in Figure 3C). Indeed, the effects of both sets of mismatches conformed to this simple model. Thus, our results support a model in which mismatches between the DNA and crRNA slow R-loop propagation and have no additional effects on the cleavage rates of either DNA strand.

**Figure 3.**
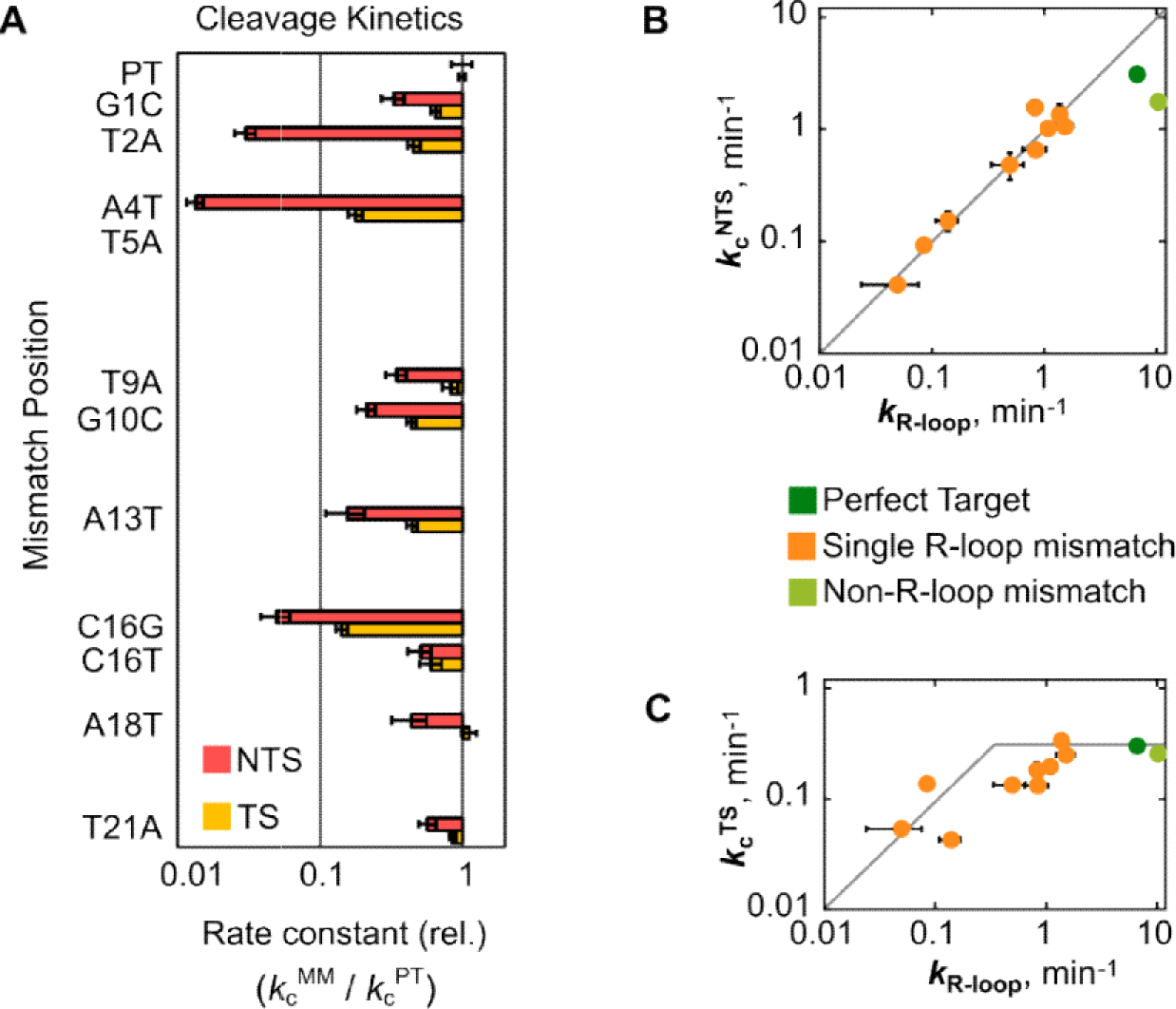
Mismatches impede R-loop propagation, affecting binding and cleavage. (A) Bar graphs represent the rates of cleavage for a mismatched target normalized to the matched target rate. Error bars reflect the normalized standard error of the fit. Red bars: NTS cleavage; Yellow bars: TS cleavage. See Also Figure S3 and Table S2. (B, C) Plots showing the correlation between effects of mismatches on binding (R-loop formation) and DNA cleavage. As the rate of R-loop formation decreases from mismatches, the rate of NTS cleavage mirrors this decrease (panel B). Values of *k*_R-loop_ are the *k*_max_ values derived from Cas12a binding fits. The grey line denotes a 1:1 correspondence between the rate constants for R-loop formation and NTS cleavage. In contrast, the observed rate of TS cleavage is not decreased by mutation unless the rate of R-loop formation becomes slower than the rate of cleavage of the matched target (panel C). The gray line, starting from the right side of the graph with a value of 0.33 min^−1^ (*k*_c_^TS^, perfect target), remains flat until reaching 0.33 min ^−1^, where it descends in parallel with the rate of R-loop formation.

### Cas12a binding remains rate-limiting at a physiological Mg^2+^ concentration

Biological Mg^2+^ concentrations are between 0.2 mM and 1 mM in mammalian nuclei (Gunther, 2006) and only slightly higher in bacterial cells (Lusk et al., 1968), much lower than the 5-10 mM used in most *in vitro* studies. To extend our kinetic framework to a cellular milieu, we also conducted reactions at the lower Mg^2+^ concentration of 1 mM (Figure S4). Cas12a binding to its perfect target DNA was slowed by 30%, to 4.9 (± 0.2) × 10^9^ M^−1^ min^−1^, and Cas12a(D908A) dissociation was accelerated 6-fold (to 0.0020 ± 0.0004 min^−1^). Cleavage of the NTS, with a *k*_c_^NTS^ value of 1.2 ± 0.1 min^−1^, remained orders of magnitude faster than Cas12a dissociation. Thus, at a near-physiological Mg^2+^ concentration, Cas12a remains an efficient DNA nuclease. The increased dissociation rate demonstrates the importance of Mg^2+^ for stability of the ternary complex, but the affinity remains very high (*K*_d_ = 410 fM), and DNA binding is still rate limiting for cleavage. Thus, it is likely that DNA binding remains rate limiting for cleavage *in vivo* and that target specificity is determined by differences in the rate constant for Cas12a binding, not differences in affinity.

## DISCUSSION

Cas12a is currently a subject of intense interest for its simplicity and potential for high target specificity *in vivo*. Here, we provide a detailed kinetic mechanism that outlines a mechanistic basis for its extraordinary specificity. A quantitative kinetic analysis of the physical and chemical steps leading up to cleavage of the two target strands shows that binding of Cas12a to target DNA involves two kinetically separable steps (Figure 4). An initial step likely represents PAM recognition, which is rapid and reversible. Next, the R-loop forms with a maximal rate constant of ~6 min^−1^, as revealed by a plateau in the binding rate constant and by 2-AP fluorescence experiments. The same rate constant is obtained whether the 2-AP probe is near the PAM-proximal end or the PAM-distal end, indicating that the R-loop propagates quickly after initiating adjacent to the PAM. After R-loop formation, the NTS is cleaved rapidly, such that R-loop formation limits the cleavage rate under saturating conditions. Following this cleavage event, there is additional trimming of four nucleotides of the NTS and cleavage of the TS 10-fold more slowly. The DNA cleavage steps are not significantly impacted by single mismatches between the crRNA and the target DNA. Instead, the decreased observed cleavage rates arise from decreases in the rate of R-loop formation.

Surprisingly, DNA cleavage is less precise than previously thought, as the NTS is trimmed towards the PAM and the TS is cleaved at multiple positions. These cleavage events imply that there is considerable mobility of the NTS and the RuvC domain and that there is flexibility in positioning of the TS in the nuclease active site. Biologically, this trimming is significant because it leads to non-cohesive DNA ends, which require filling in by a polymerase but could also be subjected to resection during DNA repair (Chang et al., 2017). Additionally, the expected rapid release of the PAM-distal cleavage product following TS cleavage (Singh et al., 2017) would produce an unprotected 5´-overhang available for binding by repair factors or annealing to a new DNA with microhomology.

**Figure 4.**
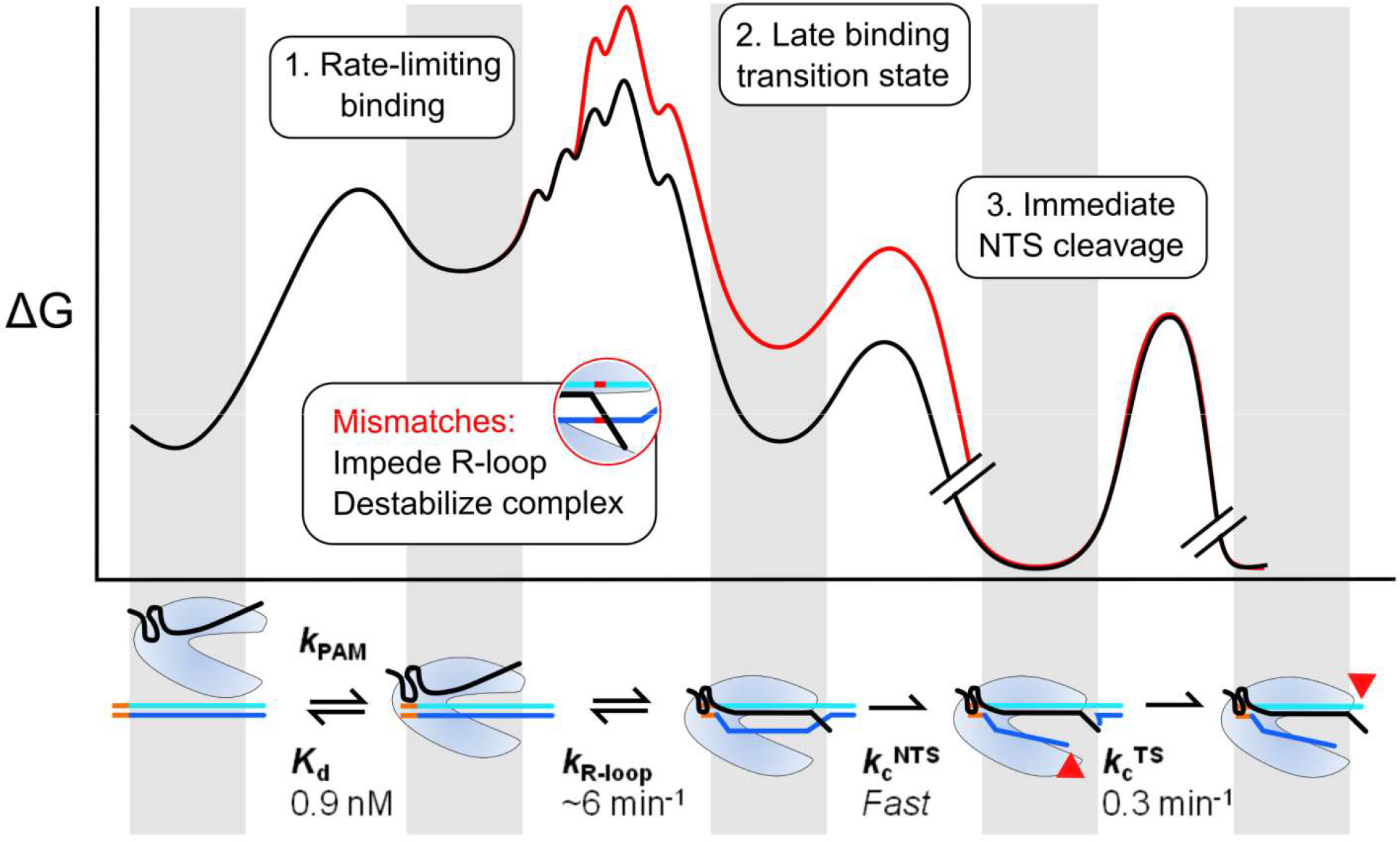
Model and kinetic framework for Cas12a target recognition and cleavage. Cas12a specificity is determined during R-loop formation. Due to the late transition state for R-loop formation, Cas12a is able to discriminate against mismatches across the R-loop. Cleavage of the NTS strand occurs rapidly, such that R-loop formation is rate-limiting for cleavage even under saturating conditions. TS cleavage occurs ~10-fold more slowly. A mismatch (red curve) increases the energy barrier for R-loop formation, resulting in a decreased rate of R-loop formation with no direct effect on the DNA cleavage steps. Breaks in the free energy profile note irreversible cleavage steps. Valleys correspond to the ground states of Cas12a shown in the cartoon below the profile (aligned by grey panels).

Our kinetic analysis showed that substrate dissociation is orders of magnitude slower than DNA cleavage. This result indicates that DNA binding is rate limiting for cleavage under subsaturating conditions, and therefore DNA binding controls the level of target specificity. Importantly, this behavior is robust, with the same regime holding at low, physiological Mg^2+^ concentration and for targets with mismatches. Despite DNA binding being rate-limiting for DNA cleavage, substantial penalties were observed for mismatches throughout the R-loop. This surprising result implies a late transition state for R-loop formation, with R-loop propagation being readily reversible until most of the 20 base pairs have formed. The late transition state may arise as a consequence of the concurrent displacement of the non-target DNA strand, allowing a reversible ‘exchange’ reaction between the DNA duplex and the R-loop that may be augmented by features of Cas12a. Alternatively or in addition, features of the contacts of the R-loop and/or the NTS with Cas12a may contribute to the reversibility of R-loop formation.

Our finding that Cas12a binding to target DNA is rate-limiting for cleavage suggests that the levels of discrimination against off-target cleavage in cells are determined by the extent to which the rate of target binding, not the affinity, is decreased by a given mismatch (Bisaria et al., 2017). Indeed, the reported patterns of off-target DNA cleavage *in vivo* show decreased discrimination at the PAM-distal end and in the center of the R-loop (most notably for positions 9 and 10) (Kim et al., 2016; Kim et al., 2017; Kleinstiver et al., 2016), closely resembling the effects of mismatches on the binding rate constant rather than effects on equilibrium binding (see Figure 2).

Our results also highlight a key difference between Cas12a and Cas9. Unlike Cas12a, the positions of strong discrimination against mismatches are confined to a PAM-proximal ‘seed’ region for Cas9, as measured both in cells by DNA cleavage and *in vitro* by binding experiments (Boyle et al., 2017; Hsu et al., 2013; Singh et al., 2016; Sternberg et al., 2014). Analogous to our results for Cas12a and supported by recent kinetics experiments on Cas9 (Gong et al., 2018), it appears likely that DNA binding also limits the rate of cleavage by Cas9 *in vivo*. Thus, the limited seed region for Cas9 specificity implies that the transition state for R-loop formation is reached upon formation of fewer base pairs than for Cas12a. We conclude from this difference that the position of the transition state for binding can be influenced by properties of the Cas endonuclease, not just by the intrinsic reversibility of the R-loop, which would be equivalent for Cas12a and Cas9. Further, the variation in transition state position suggests the possibility of engineering nucleases to shift the transition state even further into the PAM-distal region of the R-loop, extending the region and the overall levels of specificity for gene editing applications.

## ACKNOWLEDGEMENTS

We thank Prof. Ken Johnson and Shanzhong Gong for use of the stopped-flow fluorimeter and for guidance with the 2-AP fluorescence experiments, Prof. Ailong Ke for providing a Twin-Strep-Sumo expression vector, and members of the Russell and Finkelstein labs for helpful discussions and comments on the manuscript. This work was supported by NIGMS grants P01GM066275 (to D. Herschlag, R.R. Co-I) and R01GM124141 (to I.J.F.) and by Welch Foundation grants F-1563 (to R.R.) and F-1808 to (I.J.F.).

## AUTHOR CONTRIBUTIONS

Conceptualization, all authors; Methodology, I.S., F.M., I.J.F., and R.R.; Investigation, I.S. and F.S.; Writing - Original Draft, I.S. and R.R.; Writing - Review & Editing, all authors; Funding Acquisition, I.J.F. and R.R.

## STAR METHODS

### Cas12a Cloning and Purification

An *E. coli* codon-optimized gene encoding AsCas12a was purchased from BioMatik. The gene was cloned into a custom pET-19 based expression vector pIF191 (WT) and pIF255 (D908A) and transformed into BL21 star (DE3) cells (ThermoFisher). Cas12a contained a His_6_/Twin-Strep/SUMO N-terminal fusion. Single colonies were used to inoculate Terrific Broth supplemented with 50 μg/mL carbenicillin at 37 °C. An aliquot of the starter culture (4 mL) was then used to inoculate 1 L of Terrific Broth. When the culture reached an OD_600_ of ~0.6, 1 mM IPTG was added and cultures were further incubated at 18 °C for 24 hrs. Cells were then lysed in 20 mM Na-HEPES, pH 8.0, 1 M NaCl, 1 mM EDTA, 5% glycerol, 0.1% Tween-20, 1 mM PMSF, 2000 U DNase (GoldBio), and 1X HALT Protease Inhibitor (ThermoFisher) and homogenized. The lysate was clarified by centrifugation and was applied to a hand-packed Strep-Tactin Superflow gravity column (IBA Life Sciences). Cas12a was eluted with 20 mM Na-HEPES, 1 M NaCl, 10 mM desthiobiotin, and 5 mM MgCl_2_. SUMO protease (2.4 μM) was added to the eluate and the solution was dialyzed into 500 mM NaCl overnight at 4 °C. The sample was then fractionated over a HiLoad 16/600 Superdex 200 Column (GE Healthcare) equilibrated in 20 mM Na-HEPES, pH 8.0, 500 mM NaCl, 5 mM MgCl_2_, and 2 mM dithiothreitol (DTT). Fractions containing Cas12a were determined by SDS-PAGE, pooled and concentrated to ~10 μM using a 30 kDa centrifuge concentrator (Millipore) (See Figure S1C). Small aliquots were flash frozen in liquid nitrogen and stored at −80 °C.

### DNA Substrate Preparation

Oligonucleotides corresponding to the Target Strand (TS) and Non-target Strand (NTS) (Integrated DNA Technologies) (see Table S1) were 5´-radiolabeled with [γ-^32^P]ATP (Perkin-Elmer) using T4 polynucleotide kinase (New England Biolabs). Radiolabeled oligonucleotides were purified by polyacrylamide gel electrophoresis, eluted in TE buffer, and stored at −20 °C. Target DNA duplexes were formed by incubating either the radiolabeled TS or NTS (25 nM) with four-fold excess of the unlabeled complementary strand in annealing buffer (50 mM Na-MOPS pH 7.0, 100 mM NaCl). The mixture was heated at 90 °C for 5 min, slow-cooled to 25 °C, and diluted to ~1 nM duplex (corresponding to the calculated concentration of the radiolabeled strand).

### Cas12a-crRNA Assembly

The Cas12a-crRNA complex was assembled for experiments each day using purified Cas12a and a minimal precursor crRNA (pre-crRNA) that had a single uridine upstream of the Cas12a cleavage site (purchased from IDT). The crRNA precursor sequence is: 5´-UAAUUUCUACUCUUGUAGAUGUGAUAAGUGGAAUGCCAUGUGGA. Assembly reactions were performed by incubating Cas12a with a 2-fold excess of the precursor crRNA for 25 min at 25 °C in 50 mM Na-MOPS, pH 7.0, 120 mM NaCl, 5 mM MgCl_2_, 2 mM DTT, and 0.2 mg/ml bovine serum albumin. The assembled complex was then diluted, maintaining the same buffer conditions, and stored on ice until use.

### Measurement of Cas12a binding kinetics

Reactions were performed in the same buffer conditions as the assembly reaction. DNA binding was initiated by the addition of a trace amount of target DNA duplex, radiolabeled on the NTS, to the assembled Cas12a-crRNA complex. At various time points, 2 μl of the 20 μl binding reactions were withdrawn and added to 4 μl of an ice-cold chase solution consisting of 3X loading dye (15% glycerol, 5 mM Tris-Cl, pH 8.0, 0.015% xylene cyanol) supplemented with 50 mM Na-MOPS, pH 7.0, 120 mM NaCl, 20 mM EDTA, and 100 nM unlabeled perfect target duplex. Samples were electrophoresed on a 12% native gel at 4 °C, exposed to a phosphorimager screen overnight, and then scanned using a Typhoon FLA 9500 (GE Healthcare). Bands of radioactivity corresponding to unbound and bound DNA complexes were quantified using ImageQuant 5.2 (GE Healthcare). For experiments performed at 1 mM Mg^2+^, Cas12a was first assembled with the crRNA under standard conditions at 5 mM Mg^2+^ and then diluted to 1 mM Mg^2+^ prior to initiation of reactions.

### Measurement of Cas12a dissociation kinetics

A DNase-inactive mutant (Cas12a(D908A)) was used for dissociation reactions in the presence of Mg^2+^. Purified Cas12a(D908A) was loaded with the minimal pre-crRNA as described above. Cas12a(D908A)-crRNA was then mixed with a trace amount of target DNA duplex labeled on the NTS for at least one hr at 25 °C to allow complete binding of the duplex. For mismatch targets that bound slowly, binding incubation periods were increased to 2 hrs. Dissociation reactions were performed at 10 nM Cas12a(D908A) and initiated by adding the unlabeled perfect target DNA in 50-fold excess over the protein concentration to ensure no further binding or rebinding of labeled DNA during the time course. Aliquots (2 μl) were withdrawn over the course of ~24 hrs, mixed with 2 μl of ice-cold 6X loading dye (30% glycerol, 10 mM Tris-Cl, pH 8.0, 0.03% xylene cyanol), and stored at 4 °C. Samples were electrophoresed on a 12% native gel at 4 °C, imaged with a phosphorimager, and analyzed as described above. For experiments performed at 1 mM Mg^2+^ or in the absence of Mg^2+^, Cas12a (wild-type or D908A) was assembled with the crRNA as described above and then diluted to 1 mM Mg^2+^ during the DNA loading step or chelated with 10 mM EDTA.

### Measurement of DNA strand cleavage rates

DNA duplexes were radiolabeled at the 5´ end of either the TS or NTS. Cleavage reactions were performed using 2, 10, and 50 nM Cas12a (pre-assembled with crRNA) and were initiated by addition of a trace amount of labeled target DNA. At various times, 2 μl samples were quenched in 4 μl of denaturing quench (90% formamide, 20 mM EDTA, 0.01% bromophenol blue, and 0.005% xylene cyanol). Our denaturing quench also included 300 nM of an oligonucleotide with no complementarity to our target sequences to reduce retention on microfuge tube walls and in the gel wells. Samples were heated to 95 °C for 3 min and immediately loaded onto a 20% denaturing polyacrylamide gel containing 7 M urea. Gels were exposed and imaged as described above. Bands corresponding to cleaved and uncleaved DNA substrate were quantified. For experiments performed at 1 mM Mg^2+^, Cas12a was first assembled with the crRNA under standard conditions at 5 mM Mg^2+^ and then diluted to 1 mM Mg^2+^ when it was transferred into the cleavage reactions.

### Measurement of R-loop propagation rate

Non-target strands were purchased as oligonucleotides with 2-aminopurine substituted at position 4 or 18 (IDT). The target duplex was formed by annealing the NTS and TS (10 μM each) in a thermocycler in 2-AP Annealing Buffer (10 mM Tris-Cl, pH 8.0, 50 mM NaCl, 1 mM EDTA). The duplex was then diluted to 600 nM with Running Buffer (100 mM NaCl, 50 mM Tris pH 8.0, 5 mM MgCl_2_, 2 mM DTT) in an amber tube. Cas12a was assembled with a 2-fold excess of precursor crRNA for 25 min at room temperature in binding buffer (50 mM Tris-Cl, pH 8.0, 100 mM NaCl, 5 mM MgCl_2_, 2 mM DTT). The SF 2004 series stopped-flow (KinTek Corp.) ports were washed with 600 μl (port volume) of 2 N NaOH, 2 N HCl and 5 × 600 μl of deionized water before experiments. For each experiment, 300 μl of the target DNA was loaded into port A and 300 μl of 1 μM Cas12a solution was loaded into port B. Replicate data sets were combined and analyzed using the KinTek Global Kinetic Explorer to obtain rates.

